# A Model of Mitochondria in the Rat Hepatocyte

**DOI:** 10.1101/2021.12.17.473135

**Authors:** William Bell, Anita T. Layton

## Abstract

Mitochondria are a key player in several kinds of tissue injury, and are even the ultimate cause of certain diseases. In this work we introduce a new model of mitochondrial ATP generation in liver hepatocytes of the rat. Ischemia-reperfusion is an intriguing example of a non-equilibrium behaviour driven by a change in tissue oxygen tension. Ischemia involves prolonged hypoxia, followed by the sudden return of oxygen during reperfusion. During reperfusion, we predict that the build up of succinate causes the electron transport chain in the liver to temporarily be in a highly reduced state. This can lead to the production of reactive oxygen species. We accurately predict the timescale on which the electron transport chain is left in a reduced state, and we observe levels of reduction likely to lead to reactive oxygen species production. Aside from the above, we predict thresholds for ATP depletion from hypoxia, and we predict the consequences for oxygen consumption of uncoupling.

## 1 Introduction

Mitochondria are crucial to understanding the pathophysiology of the liver in a wide array of circumstances. Some of the ways in which liver pathophysiology specifically is impacted by or mediated through mitochondria are not unique to the liver. Hypoxia is a common consequence of liver disease [55] and ischemia-reperfusion injury is a common consequence of liver transplantation [41] but they are certainly not unique to the liver. Other mitochondria-related pathophysiologies are unique to the liver. For instance alcoholic fatty liver disease is of course unique to the liver, and mitochondria are causally-connected to its development [61]. These unique problems for the liver also are associated with special adaptive responses. For instance during alcohol consumption, in healthy rats there is a large increase in liver oxygen tension [47]. This could potentially prevent hypoxia but could also increase oxidative stress, which is associated with alcohol-related liver injury [61]. In light of all these conditions, understanding mitochondria is crucial to understanding liver pathophysiology, but it is also necessary that the liver is recognized as the unique organ that it is when investigating mitochondria-related liver pathophysiology.

Since our interest is in a cellular organelle, we take a subcellular modelling approach to study this system. We study a system of ordinary differential equations which describe pyruvate oxidation, the tricarboxylic acid (TCA) cycle, and oxidative phosphorylation (OX-PHOS). We take a model originally developed for cardiac muscle tissue [60], and since adapted to study the proximal tubule in the kidney [14], and adapt it into a model for hepatocytes. We opt to preferentially study hepatocytes within the liver because by volume the liver is composed of 80% hepatocytes [3]. The second-most common cell type, stellate cells, make up only 5-8% of liver cells [18]. Hepatocytes are also are the basis for many critical liver functions, including those related to energy metabolism, such as glycogenesis and gluconeogenesis [42].

We aim to develop a mathematical model of the hepatocyte mitochondrial function in both healthy and pathological conditions. Under baseline (healthy) conditions, we need to consider a range of values both to account for uncertainty and to account for heterogeneity between hepatocytes [4]. For our stated purpose of studying mitochondrial function, gradients of oxygen and preferences of metabolic pathways within liver lobules are particularly important. Other quantities are highly uncertain and must be varied for that reason. In particular ATP consumption and the contribution of glycolysis to liver function (the latter is also subject to spatial heterogeneity). When we examine the sensitivity of our model under health conditions, we’ll be interested especially to identify the sensitivity of the default conditions to all the features mentioned here.

We then apply the model to investigate several causes of liver dysfunction, namely: mitochondrial disease (and especially those based on OXPHOS-mediated disorders), fatty liver disease (which especially causes oxidative stress, leading to OXPHOS dysfunction and Cytochrome C release), hypoxia, and acute alcohol consumption (which causes oxidative stress). Crucially, many of these either lead to or follow from OXPHOS dysfunction, and so our aim will be two-fold: to replicate conditions of a highly reduced OXPHOS chain, the natural preceding event to oxidative stress, and to consider OXPHOS dysfunction once it occurs.

## 2 Method

### 2.1 The Model

The model is adapted from a model for cells in the kidney of the rat. The biochemical reactions and transport kinetics in the two tissues are relatively similar, and so the model structure isn’t significantly altered for this work. The only structural change to the kidney model is the introduction of glutamate dehydrogenase activity, described below. The key changes relative to either the proximal tubule (PT) or medullary thick ascending limb (mTAL) model were to the parameters, including some clamped states of the model. This happened in two phases: first we changed parameters that had been measured experimentally, and second we fitted some less measurable parameters. Section 2.2 addresses this second fitting phase. In this section we address experimentally determined parameter values.

Several important quantities for the model can be measured directly, they are listed in Table 1. ATP consumption ranges were chosen based on the activity of Na-K-ATPase in mice [25] and an estimate in rats that 20% of ATP consumption (via a proxy of oxygen consumption) is due to Na-K-ATPase [2]. This gave us a reasonable-appearing estimate that the maximal ATP consumption was roughly half of that of a proximal tubule cell in the kidney. Proximal tubule cells have particularly large energy requirements, suggesting this is reasonable [14]. Glycolysis-based ATP production was taken to be a third of the maximal glycolysis-mediated pyruvate and lactate production found in rats in Fedatto et al. [15].

**Table 1:** A list of parameters used in our model that were based on experimental data.

| Parameter | Values for Liver | Reference |
| --- | --- | --- |
| Fractional Mitochondrial Volume (L mito/L cell) | 0.23 | [3] |
| Fractional Cytoplasmic Volume (L cyto/L cell) | 0.70 | [3] |
| Cytoplasm Water Fraction (L water/L cyto) | 0.77 | [53] |
| Mitochondria Water Fraction (L water/L mito) | 0.58 | [54] |
| Matrix NAD+NADH concentration (mmol/L matrix) | $9.4 \times 10^{-3}$ | [8] |
| Matrix concentration of co-enzyme Q (mmol/L matrix) | $6.49 \times 10^{-3}$ | [5] |
| Intermembrane concentration of Cytochrome C (mmol/L IM) | $4.39 \times 10^{-3}$ | [5] |
| Protein Density of Mitochondria (mg prot/L mito) | $6.45 \times 10^5$ | [29] |
| Liver-to-heart Ratio Complex I | 1.2 | [5] |
| Liver-to-heart Ratio Complex III | 0.525 | [5] |
| Liver-to-heart Ratio Complex IV | 0.40 | [5] |
| Mitochondrial Oxygen Tension (mmHg) | 35 | [39] |
| Cytosolic Potassium Concentration (mmol/L cyto) | 88 | [57] |

The model structure was altered to include one additional flux, corresponding to the activity of glutamate dehydrogenase, which in mammalian cells converts glutamate into alphaketogluterate, a Krebs cycle intermediate. This flux was represented in our model with simplified kinetics as follows:

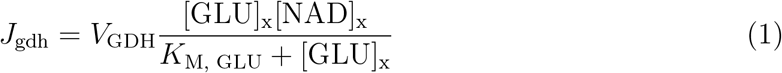

The activity of glutamate dehydrogenase is a source of alphaketogluterate and NADH in the mitochondrial matrix, and consumes glutamate and NAD^+^. *V*_GDH_ and *K*_M, GLU_ were chosen based on experimental measurements [27]. That paper uses a model of the kinetics that was even simpler, not including NAD^+^, for that reason, we treat that paper as measuring 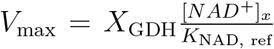, where *K*_NAD, ref_ is a baseline concentration of NAD^+^ present in the mitochondrial matrix. We use physiologically reasonable concentrations to choose *K*_NAD, ref_, assume for calculation that this was the NAD^+^ concentration present in the mitochondrial matrix in the work of Jonker et al. [27], and then *V*_GDH_ may be calculated simply as 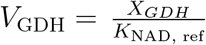. Our model requires us to model the dependence on NAD^+^ concentration because otherwise the model may consume excessive NAD^+^.

### 2.2 Model Parameter Fitting

With a minor increase (16%) in the hydrogen leak permeability relative to the proximal tubule, we are able to find a suitable fit for the State 2 and 3 oxygen consumption as measured in isolated liver mitochondria [21]. In that work they measure the State 2 measurements of oxygen consumption to be 0.30 mM/s, and the State 3 measurements of oligomycin-sensitive (or Complex IV-dependent) oxygen consumption to be 2.0 mM/s. With adjustments to the hydrogen leak we get a good fit for the State 2 and 3 oxygen consumption. The results of the fit are recorded in Table 2.

**Table 2:**
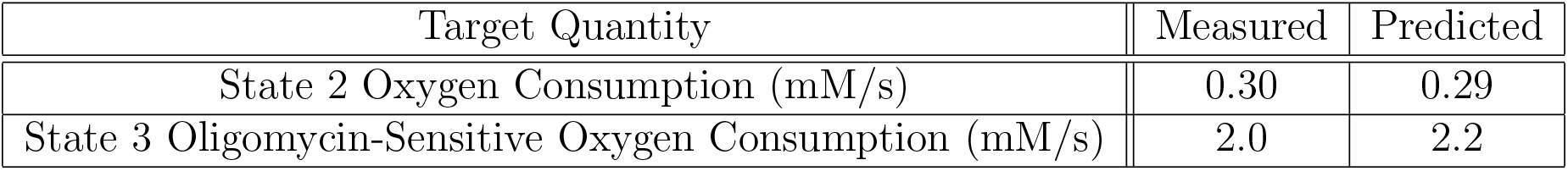
Values used to fit unknown parameters and the predicted values.

### 2.3 Simulations

While there is a variety of mitochondria-mediated liver pathologies, they tend to progress through OXPHOS dysfunction or oxidative stress. The latter is attenuated by uncoupling effects, or hydrogen leak permeability. Uncoupling reduces the effect of oxidative stress because when the proton motive force is less strong, OXPHOS complexes are better able to transfer their electrons through the electron transport chain instead of staying in a reduced state that increases the risk of free radical formation. Oxidative stress can also initiate mitochondrial permeability transition (mPT), which may reduce the concentration of cytochrome C in the intermembrane space. Oxidative stress can also produce OXPHOS dysfunction.

OXPHOS dysfunction is represented by some combination of reductions in the activities of Complex I, III, IV, and ATP Synthase (*X_CI_*, *X_CIII_*, *X_CIV_*, and *X_F1_*). We consider combinations in reductions of these activities because many OXPHOS dysfunctions are not specific to one OXPHOS enzyme [37]. We consider successive reductions in the activity of each OXPHOS enzyme by 25% to as low as a quarter of the baseline enzyme activity.

Oxidative stress is frequently a consequence of the cell being in an energized state where the redox state of the cell’s electron transport chain is too reduced. We consider two mechanisms that are able to produce oxidative stress: high glycolysis (four times the default flux) or via reperfusion following a period of ischemia (ischemia involving zero oxygen tension, smaller adenine nucleotide and nicotinamide pools). In the latter case, we consider the consequences of total hypoxia (no oxygen) over a period of 50,000 seconds (or roughly 14 hours) with a nicotinamide pool (the combined concentrations of NADH and NAD^+^) that is half the normal size, and a pool of cytosolic adenine nucleotides (the combined concentrations of ATP, ADP, and AMP) that is a quarter of normal levels [28, 58]. Following this period of ischemia, liver oxygen tension returns to normal but other features of the cell do not. We then recorded the reperfusion behaviour until it appeared to settle at a new baseline.

Aside from the above cases, we consider uncoupling on its own and hypoxia. The latter is a known consequence of liver disease, and especially alcoholic liver disease in rats [47, 55]. We considered oxygen tensions reduced by tenths from the default oxygen tension of 36 mmHg, to a low of 3.6 mmHg. We also consider an oxygen tension of 0.5 mmHg to gauge the effects of more extreme hypoxia. For uncoupling, we consider 2x, 5x, and 10x typical hydrogen leak permeability (P_H,leak_).

## 3 Results

### 3.1 Baselines

Key model predictions are shown in Table 3. Under baseline conditions, the liver model predicts a cytosolic ATP concentration of 6.81 mM, an energy charge of 0.97, and an ATP/ADP ratio of 14.55. These compare favourably with at least some experimental estimates. For instance Schwenke et al. measure the cytosolic ATP concentration to be 6.18 mM. The cytosolic ATP/ADP ratio was found to be 6.94 in the same study [49], although the ATP/ADP ratio varies widely between preparations [49, 62]. Other predicted measures of the energetic state of the liver also appear to accord well with at least some experimental results. For instance the electrical potential gradient across the inner membrane of liver mitochondria was predicted to be 165 mV (physiological conditions treated as a positive quantity). This is within experimentally observed ranges. Porter and Brand observe an electrical potential gradient of 149 mV [46]. Other studies on isolated hepatocytes have found the electrical potential gradient to be 154 mV in young rats [20]. Some studies report even lower values for the electrical potential gradient such as 108 mV [52] or 118 mV [40], both in perfused liver. Other studies working with isolated rat liver mitochondria have reported electric potential gradients as high as 188 mV [45]. In the above cases, the proton motive force was also frequently measured. There is a wide range of experimental measurements, which encompasses our predictions, we predicted the proton motive force to be 168 mV. Experiments have found the proton motive force to range from 125 mV [52] to 191 mV [20].

**Table 3:** The model under typical conditions and various relevant deviations from normal conditions.

|  | O<sub>2</sub> Con-<br>sumption<br>(nmol O<sub>2</sub><br>mg<sup>-1</sup> s<sup>-1</sup>) | ATP<br>Generation<br>(nmol<br>ATP mg<sup>-1</sup><br>s<sup>-1</sup>) | P/O | Proton<br>Motive<br>Force<br>(PMF,<br>mV) | [ATP]<sub>c</sub><br>(mM) | [ATP]<sub>c</sub><br>/[ADP]<sub>c</sub> |
| --- | --- | --- | --- | --- | --- | --- |
| **Base Case** |  |  |  |  |  |  |
| P<sub>O2</sub> = 36mmHg<br>pH<sub>c</sub> = 7.20 | 0.59 | 2.04 | 1.72 | 168.3 | 6.81 | 14.55 |
| **P<sub>O2</sub> Variation** |  |  |  |  |  |  |
| P<sub>O2</sub> = 50mmHg | 0.59 | 2.04 | 1.71 | 169.49 | 6.82 | 14.88 |
| P<sub>O2</sub> = 12mmHg | 0.59 | 2.04 | 1.72 | 167.67 | 6.8 | 14.30 |
| P<sub>O2</sub> = 8mmHg | 0.59 | 2.04 | 1.73 | 166.56 | 6.79 | 13.87 |
| **Q<sub>ATP,max</sub> Variation** |  |  |  |  |  |  |
| Q<sub>ATP,max</sub> x 0.75 | 0.43 | 1.27 | 1.5 | 169.86 | 6.86 | 16.09 |
| Q<sub>ATP,max</sub> x 1.25 | 0.76 | 2.8 | 1.84 | 168.32 | 6.76 | 13.19 |
| Q<sub>ATP,max</sub> x 1.5 | 0.93 | 3.56 | 1.92 | 167.71 | 6.68 | 11.43 |
| **pH<sub>c</sub> Variations** |  |  |  |  |  |  |
| pH<sub>c</sub> = 7.40 | 0.55 | 2.03 | 1.84 | 182.01 | 6.57 | 9.68 |
| pH<sub>c</sub> = 7.00 | 0.65 | 2.04 | 1.57 | 155.46 | 6.97 | 21.72 |
| pH<sub>c</sub> = 6.80 | 0.72 | 2.01 | 1.39 | 141.79 | 7.07 | 30.95 |
| **[K<sup>+</sup>]<sub>c</sub> Variations** |  |  |  |  |  |  |
| [K<sup>+</sup>]<sub>c</sub> = 60mM | 0.6 | 2.04 | 1.69 | 169.09 | 6.85 | 15.83 |
| [K<sup>+</sup>]<sub>c</sub> = 140mM | 0.59 | 2.03 | 1.73 | 167.82 | 6.74 | 12.62 |
| **[Mg<sup>2+</sup>]<sub>c</sub> Variations** |  |  |  |  |  |  |
| [Mg<sup>2+</sup>]<sub>c</sub> = 0.2mM | 0.59 | 2.03 | 1.72 | 169.01 | 6.7 | 11.56 |
| [Mg<sup>2+</sup>]<sub>c</sub> = 0.8mM | 0.6 | 2.05 | 1.72 | 168.98 | 6.94 | 20.01 |
| **Variations in H<sup>+</sup>**<br>**and K<sup>+</sup> leak**<br>**permeability** |  |  |  |  |  |  |
| No leaks | 0.47 | 2.09 | 2.25 | 170.38 | 6.84 | 15.47 |
| No H<sup>+</sup> leak | 0.52 | 2.07 | 2.0 | 169.69 | 6.82 | 14.95 |
| No H<sup>+</sup> leak,<br>10x P<sub>K,leak</sub> | 0.67 | 1.95 | 1.45 | 161.27 | 6.18 | 6.12 |
| No K<sup>+</sup> leak,<br>10x P<sub>H,leak</sub> | 1.29 | 1.76 | 0.68 | 165.15 | 6.78 | 13.51 |
| 10x P<sub>H,leak</sub> | 1.22 | 1.78 | 0.73 | 165.11 | 6.76 | 13.21 |
| 10x Leaks | 0.99 | 1.8 | 0.91 | 159.38 | 6.03 | 5.37 |

We calculate the P/O ratio only taking into account oxygen consumption by the mitochondria. Unlike multiple experimental studies, the P/O ratio is calculated using only mitochondrial oxygen consumption (as well with non-mitochondrial oxygen consumption) by Brand, Harper, and Taylor. Including only mitochondrial oxygen consumption, they find a P/O ratio of 1.69 [7]. This is comparable to our predicted P/O ratio of 1.72. The P/O ratio was calculated from the ATP generation, which under default conditions was found to be 2.04 nmol ATP mg^-1^prot s^-1^ by ATP Synthase, and O_2_ consumption which was found to be 0.59 nmol O_2_ mg^-1^prot s^-1^ (the P/O ratio is the ATP generation (or ATP Synthase activity) divided by 2x the oxygen consumption (or Complex IV activity)).

### 3.2 Parameter Sensitivity Analysis

Here we vary several important parameters, namely the leak permeabilities, cytosolic ion concentrations, maximal ATP consumption, and oxygen tension. For this analysis we consider large changes to these parameters and report the effects (the exact changes are noted in the first column of Table 3), rather than reporting local measures of sensitivity (see Table 3). Our sensitivity analysis indicates that only some of these perturbations have a strong impact on liver mitochondrial function, notably maximal ATP consumption and leak permeabilities.

The key parameters that most significantly alter the system’s behaviour in the ranges considered here are the maximal ATP consumption and the leak permeabilities (*P*_H, leak_ and *P*_K, leak_). Under a 1.5-fold increase in maximal ATP consumption, there’s a slightly more than 1.6-fold increase in oxygen consumption. This suggests a larger maximal ATP consumption could lead to hypoxia, due to the significantly increased oxygen consumption. However the efficiency as measured by the P/O ratio of respiration also improves with increase ATP consumption (for higher ATP consumption, the P/O ratio may be as high as 1.92). Leak permeabilities also had a major impact on the oxygen consumption. When there is no potassium leak and a ten-fold increase in hydrogen leak, the oxygen consumption more than doubles. When there is a ten-fold increase in potassium and hydrogen leak, the ATP concentration reaches its minimum, 6.03 mM, the ATP/ADP ratio hits its minimum as well of 5.37, as did the proton motive force, which gets to a minimum of 159 mV.

These results suggest that uncoupling and greater ATP demand could trigger greater oxygen consumption, possibly producing hypoxic conditions.

### 3.3 Local Sensitivity Analysis

We consider the change in state variables in hepatocytes under a change in each parameter of our model. Below we show the derivative of each state variable against each parameter, calculated using a central difference scheme with Δ*p* = 0.01*p* where *p* is the size of the parameter. The derivative is normalized relative to the baseline size of each state variable and the size of each parameter. The full local sensitivity analysis is reported in Figure 1. In Figure 2 we focus on several key variables and the parameters that are most impactful on those state variables. Hepatocytes in the liver appear to be fairly sensitive to many parameters. Several sensitivities are much higher than the comparable sensitivities in the PT and mTAL respectively. The NADH and reduced coenzyme Q concentrations were particularly sensitive to multiple key parameters. Cytosolic ATP concentrations are most sensitive to the maximal ATP consumption. The mechanism explaining the response of NADH concentrations to the activities of pyruvate dehydrogenase and alphaketogluterate dehydrogenase is the same as in the proximal tubule discussed above. Reduced coenzyme Q concentrations are strongly reactive to total NADH/NAD^+^ concentrations, and thus to pyruvate dehydrogenase and alphaketogluterate dehydrogenase activity. Aside from those effects, the reduced coenzyme Q concentration is strongly sensitive to the total available coenzyme Q, and the pool available of cytochrome C, which is negatively associated with the quantity of reduced coenzyme Q. This indicates that the concentration of reduced coenzyme Q is restricted by the total amount of coenzyme Q to a moderate extent, and that the reduction state of the coenzyme Q is also controlled by the available cytochrome C to accept electrons from coenzyme Q. Overall this indicates that the coenzyme Q and cytochrome C pools are not present in great excess of what the mitochondrion needs.

**Figure 1:**
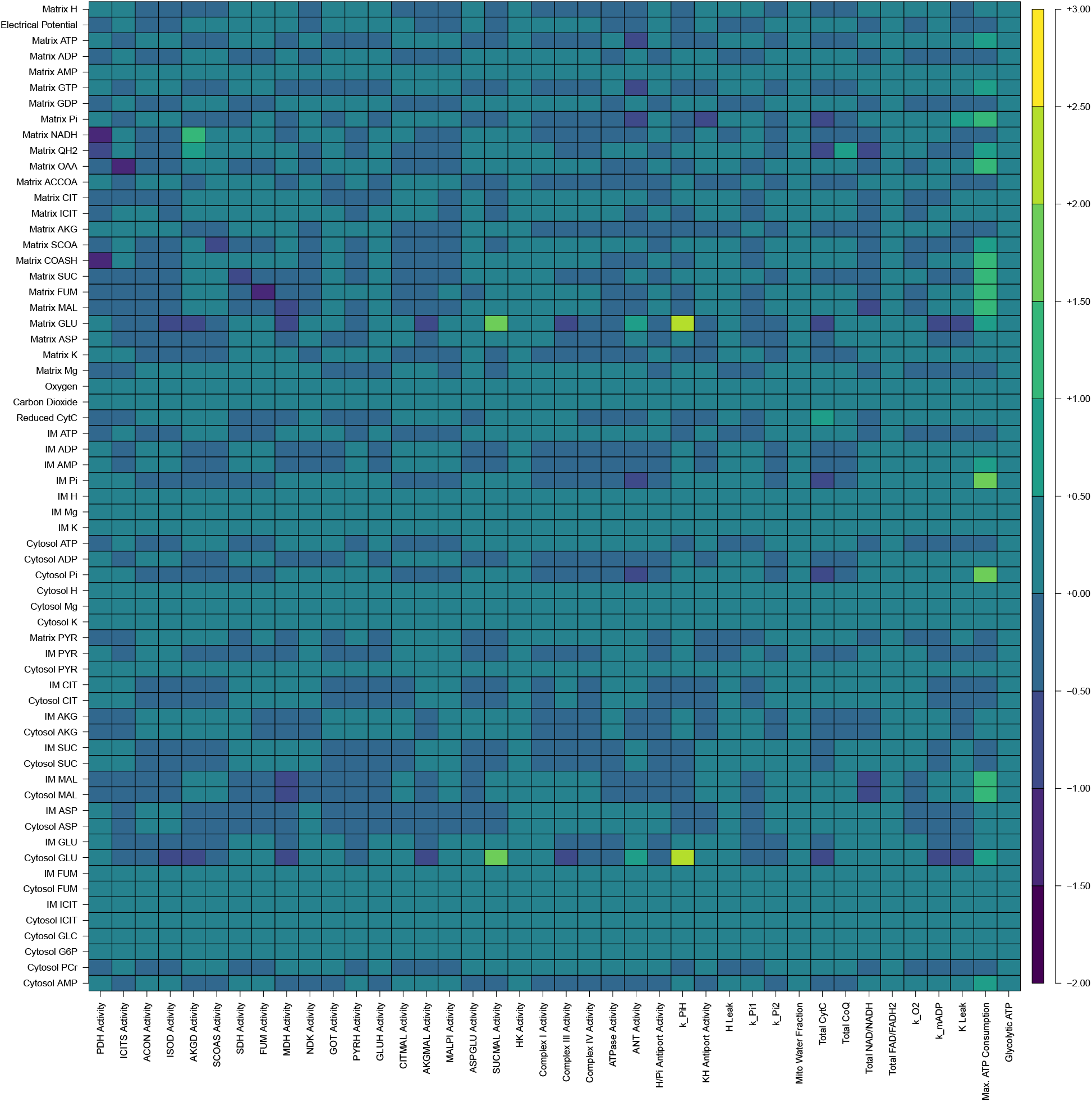
The full set of local sensitivities in the liver, for more information see section 3.3. CytC stands for cytochrome C, reduced cytochrome C refers to cytochrome C that has been donated an electron by Complex III.

**Figure 2:**
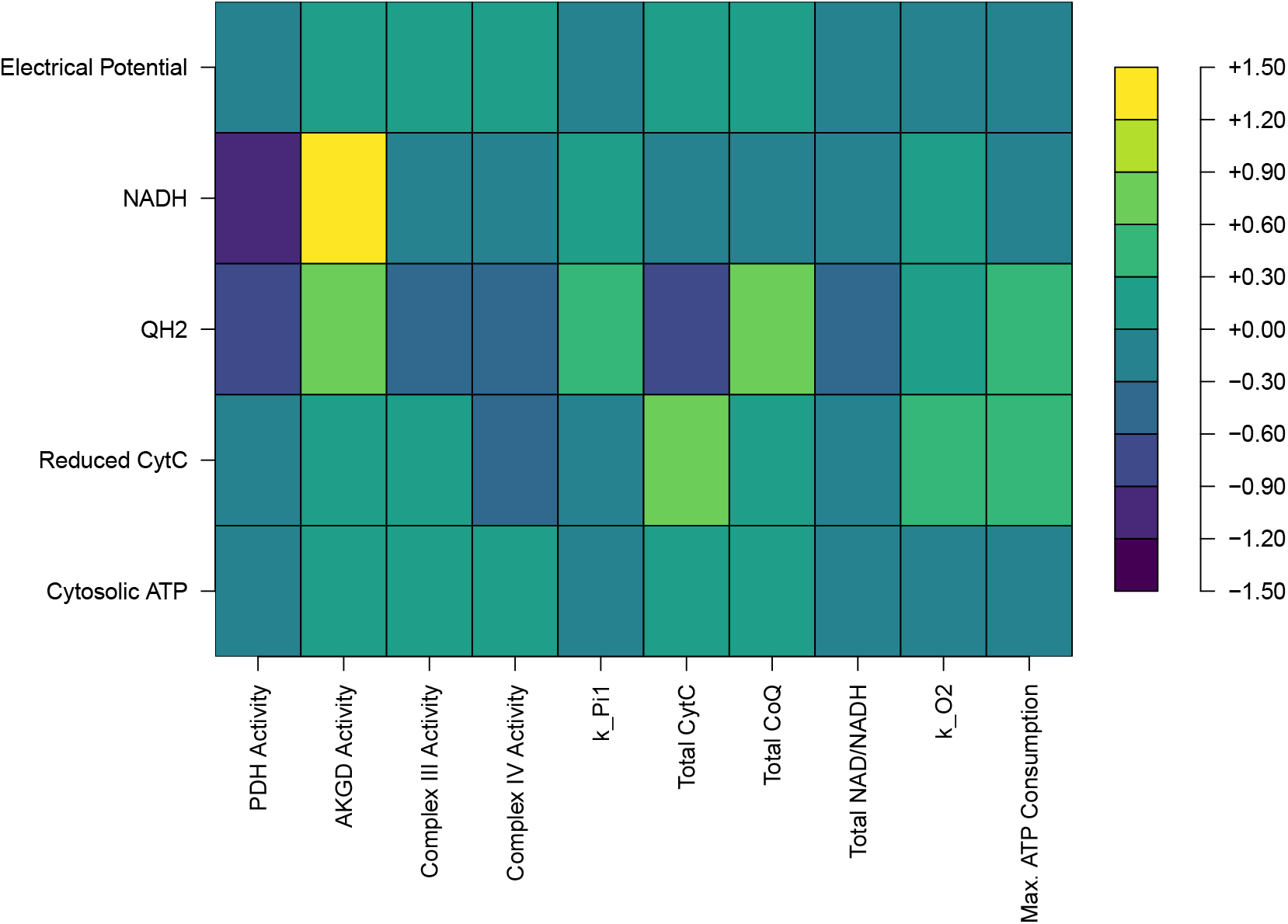
Local sensitivities of several important state variables in hepatocytes relative to certain significant parameters. CytC stands for cytochrome C, reduced cytochrome C refers to cytochrome C that has been donated an electron by Complex III. AKGD refers to alphaketogluterate dehydrogenase, PDH refers to pyruvate dehydrogenase, ANT refers to adenine nucleotide translocase, and QH_2_ is the reduced form of coenzyme Q.

### 3.4 OXPHOS Dysfunction

We describe our method for simulating dysfunction of oxidative phosphorylation (OXPHOS), that is activity of electron transport chain components and ATP synthase, above in Section 2.3. With a lowest OXPHOS complex activity of one quarter, there is little discernable consequence for cytosolic ATP consumption, and very little noticeable difference in the electrical potential gradient. With reduction to one quarter of all considered OXPHOS complex activities, the cytosolic ATP concentration differed by less than 2%. The electrical potential gradient never differed by more than 4%. Thus, impaired OXPHOS function was found to have little impact on ATP concentrations, or the electrical potential gradient of the liver. This result is consistent with the relatively few mitochondrial diseases known to affect the liver [37]. Those that do tend to be particularly severe, causing death in childhood, or progressive diseases leading ultimately to the depletion of mitochondrial DNA. The results were not significantly different under moderate hypoxia or reduced glycolysis.

Liver hepatocytes are predicted to be more robust than the proximal tubule (PT) of the kidney to OXPHOS dysfunction. The thick ascending limb (mTAL) is also more sensitive to OXPHOS dysfunction than the liver, but it has more confounding differences in tissue oxygen tension and mitochondrial volume fraction. We explore the reason for the difference between the PT and liver in robustness by considering the effect of reducing Complex III activity by 75% for hepatocyte mitochondria with PT-like parameter values for metabolic demand, glycolysis activity, or baseline OXPHOS enzyme activities (and combinations thereof). We consider another kind of robustness that the liver hepatocytes exhibit below (namely, to hypoxia). The PT has 76% of baseline cytosolic ATP concentrations under reduced Complex III activity (a difference of 0.6 mM). With no glycolysis (matching the PT) or PT-like baseline OXPHOS activities, the difference in cytosolic ATP concentration is negligible for a liver hepatocyte with Complex III dysfunction. With PT-like metabolic demand, we see 94% of baseline cytosolic ATP concentrations in hepatocytes (a difference of 0.4 mM). With both no glycolysis and PT-like metabolic demand, we see 87% of baseline cytosolic ATP concentrations (a difference of 0.9 mM). With PT-like OXPHOS activities and metabolic demand, we see 86% of baseline cytosolic ATP concentrations during Complex III dysfunction. With no glycolysis and PT-like baseline OXPHOS activities, we see negligible differences from baseline cytosolic ATP concentrations. With no glycolysis, PT-like metabolic demand, and PT-like baseline OXPHOS activities, we see 71% of baseline cytosolic ATP concentrations (a difference of 2 mM). This indicates that adjusting the glycolytic activity, OXPHOS activities, and metabolic demand to PT-like parameters is necessary and together sufficient for us to observe PT-like sensitivity to OXPHOS dysfunction.

### 3.5 Conditions producing Oxidative Stress

Oxidative stress can occur under a range of circumstances. In our simulation work, high rates of glycolysis appear to be a particularly significant potential means of producing oxidative stress. We see in Figure 3 that under these glycolytic conditions, the proton motive force is extremely high, supporting this conclusion. We consider high rates of maximal glcolysis, four times our default, similar to those observed in the presence of 20 mM of glucose [15].

**Figure 3:**
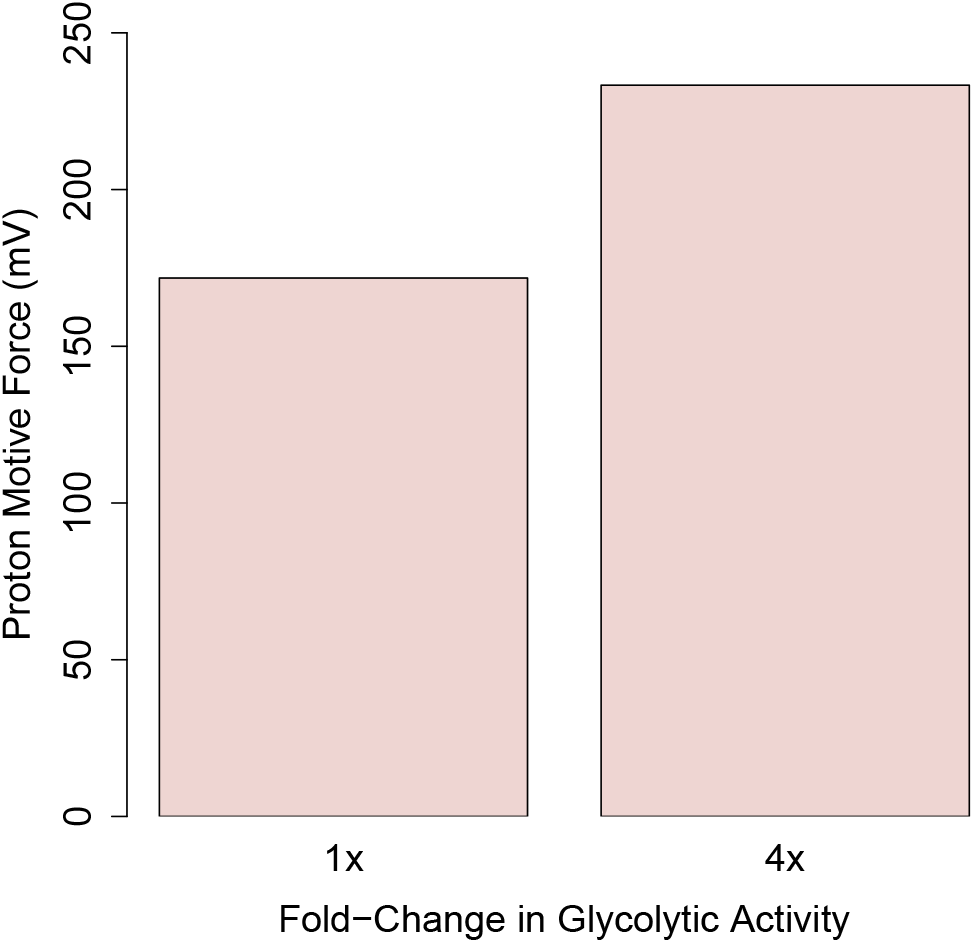
The left is the steady state proton motive force for the baseline model, and the right is the value under high rates of glycolysis (a quadrupled glycolytic activity).

### 3.6 Hypoxia

Our hypoxic simulations are outlined above in Section 2.3. The liver is only sensitive to ATP depletion under sufficiently large reductions in the oxygen tension in mitochondria. At an oxygen tension of 0.5 mmHg the cytosolic ATP concentration is reduced by 15% (to 4.3 mM). However at a tenth of normoxic oxygen tension (3.6 mmHg), cytosolic ATP concentrations are reduced by less than 2%. These results may seem inconsistent with the understood possibility of hypoxic liver injury but they are not. In both ischemia and alcohol exposure in chronic alcoholic liver disease, we see that oxygen tensions can locally get as low as can be measured quite frequently [39, 47]. Thus, it is possible that under certain kinds of disruptions of normal oxygen tension, the mitochondrial matrix becomes extremely hypoxic (below 1 mmHg oxygen tension). Mechanistically the robustness of the model to hypoxia can be explained in part by the relatively high rates of (anaerobic) glycolysis observed in liver tissue [15], this allows the tissue to continue fulfilling its ATP needs under mitochondrial dysfunction.

Comparing ATP depletion between liver hepatocytes and the kidney, we see that hepatocytes are more robust to hypoxia. We simulate hepatocytes with PT-like metabolic demand, glycolysis, and baseline OXPHOS activity in order to determine the differences that explain why they differ. We choose to compare to the PT because of the more comparable baseline oxygen tension. With 10% of baseline oxygen tension, the PT has 19% of its typical cytosolic ATP concentration (a difference of 2.1 mM). For PT-like metabolic demand, we have 73% of typical cytosolic ATP concentrations in the hypoxic hepatocyte (a difference of 1.9 mM). Without glycolysis, we have 96% of typical cytosolic ATP concentrations in the hypoxic hepatocyte (a difference of 0.3 mM). With both higher metabolic demand and no glycolysis, we have 61% of typical cytosolic ATP concentrations in the hypoxic hepatocytes (a difference of 2.7 mM). With PT-like metabolic demand, no glycolysis, and PT-like OXPHOS activity, we see 39% of typical cytosolic ATP concentrations (a difference of 4.2 mM).

### 3.7 Uncoupling

Uncoupling is used by the cell as a means of relieving oxidative stress. However uncoupling also directly reduces the P/O ratio governing the efficiency of ATP generation, and increases oxygen consumption [48, 59], which may cause hypoxia. Titrating with FCCP, a potent uncoupler, until the respiratory rate was maximal, Schönfeld et al. was able to experimentally produce a 60% increase in oxygen consumption [48]. Our model reproduces this proportional respiration rate increase with a five-fold increase in hydrogen leak. Under a two fold increase in the hydrogen leak activity, we predict a 17% increase in oxygen consumption, which in an organ with already quite significant oxygen gradients might be quite important. Already under a two-fold increase in the hydrogen leak, the P/O ratio predicted by the model to be as low as 1.32, compared to a baseline of 1.72, a 24% decrease.

### 3.8 Ischemia-Reperfusion Injury

Ischemia-reperfusion causes oxidative stress by leaving the electron transport chain in a highly reduced state temporarily. This is dangerous to the cell and the body, and can even lead to multiple organ dysfunction [43]. We wish to capture ischemia-reperfusion injury in our model and consider some candidate methods for preventing ischemia-reperfusion injury. To achieve this, we will simulate ischemia, and then we will simulate reperfusion in multiple ways. Then, we will study reperfusion in combination with interventions using pyruvate and dichloroacetate. During ischemia, the cell experiences a decline in both pyruvate and oxygen concentrations. As a consequence of ischemia, the adenine nucleotide (AMP/ADP/ATP) and nicotinamide adenine dinucleotide (NADH/NAD^+^) pools become significantly smaller. The former shrinks by three quarters [28, 58], and the latter by one half [28]. Ischemia also causes OXPHOS dysfunction that reduces the activity of Complex I, III, and IV.

For our model, we consider a shrunken pool of AMP, ADP, ATP, NADH, and NAD^+^ during reperfusion on top of restoration of typical oxygen levels after a bout of severe hypoxia (0 mmHg). We shrink the pool of adenine nucleotides by three quarters and the pool of nicotinamide adenine nucleotides by half following experimental results from Kamiike et al [28]. In this case, shown in Figure 4, we see that the NADH/NAD^+^and coenzyme Q pools are in a significantly reduced state for several minutes after reperfusion, and that the proton motive force is extremely elevated during the same period of time. This matches the timescale expected from experimental work, where it has been observed that reactive oxygen species are produced rapidly in a short period following reperfusion or reoxygenation [26]. In another trial, we consider the consequences of also including OXPHOS dysfunction. We include OXPHOS dysfunction by reducing the activity of Complex I, III, and IV by 15%, 70%, and 50% respectively following experimental results from Baniene et al. [63]. Under these circumstances, shown in Figure 5, we predictably see a somewhat more reduced coenzyme Q pool with a somewhat lower but still very high proton motive force. This is to be expected because a reduced rate of oxidative phosphorylation means that it will take longer to oxidize coenzyme Q. In other trials, we consider more moderate hypoxia (one tenth of typical) and we do not observe the same long transients. This is demonstrably due to a lack of succinate build up because when we increase the succinate levels in these trials to levels comparable to our more extreme hypoxic case, we again see long transients.

**Figure 4:**
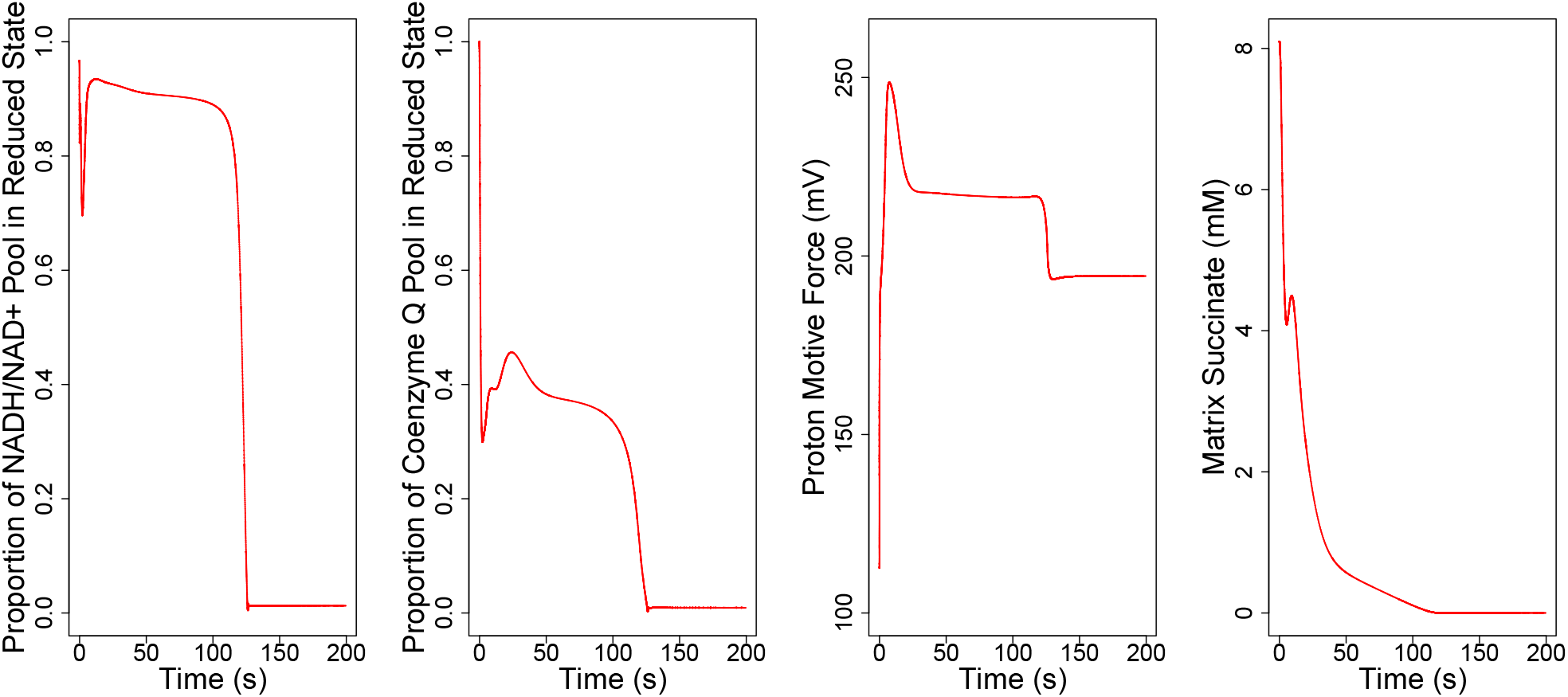
The effects of reperfusion with a smaller pool of adenine nucleotides on NADH/NAD^+^ & coenzyme Q redox state, the proton motive force, and the succinate con-centration.

**Figure 5:**
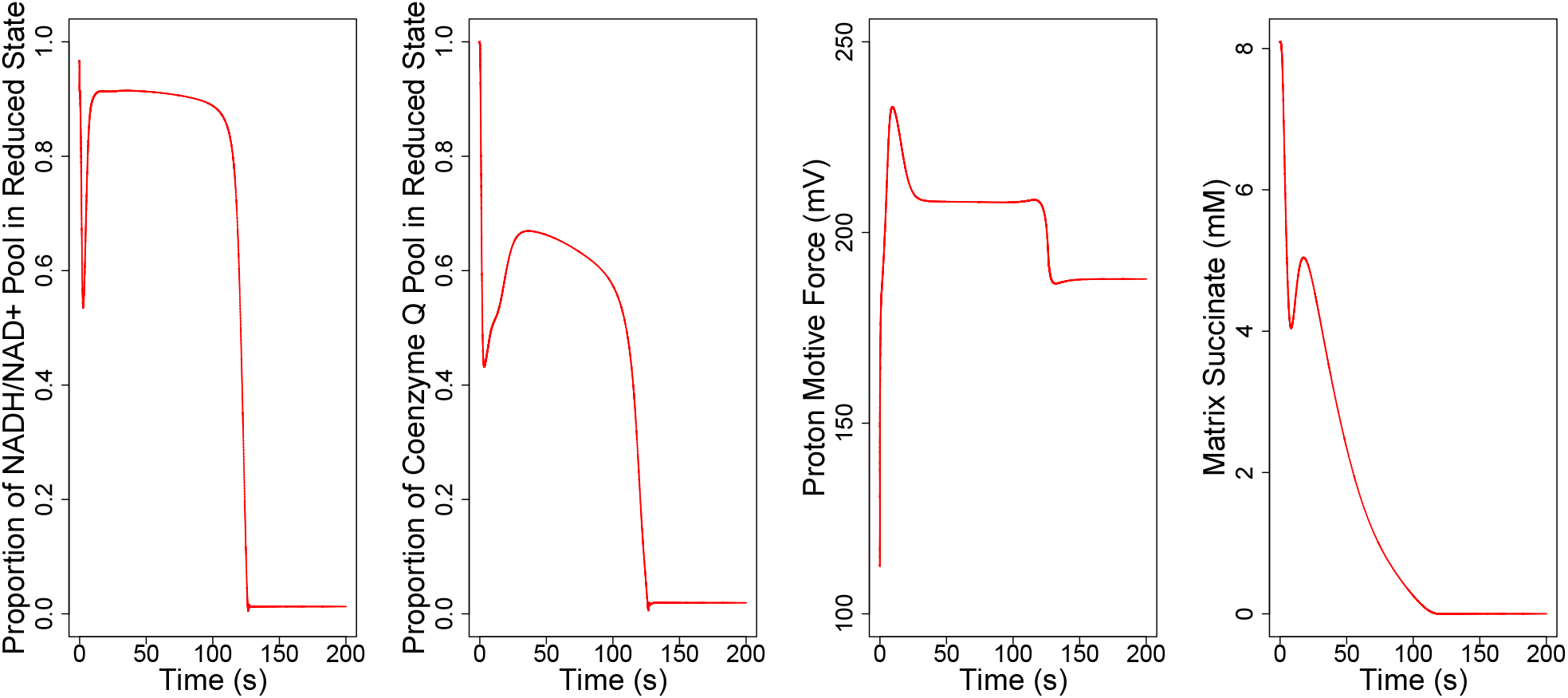
The effects of reperfusion with a smaller pool of adenine nucleotides and OXPHOS dysfunction on NADH/NAD^+^ redox state, coenzyme Q redox state, and proton motive force.

## 4 Discussion

Our model simulations tell a simple story: the liver is highly robust to ATP depletion, but possibly quite sensitive to phenomena known to cause oxidative stress. These two features of liver mitochondria happen for overlapping reasons. The liver has a high capacity for glycolytic activity [15]. This capacity means the mitochondria do not have to satisfy the hepatocyte’s metabolic burden on their own, but also can leave the mitochondria in a highly reduced state for lack of ADP to phosphorylate. This underlies our findings shown in Figure 3 that at the rates of glycolysis observed by Fedatto et al. [15] in healthy rats experiencing hyperglycemia, the electrical potential gradient is larger across the inner membrane of the mitochondria. This trade off of, on the one hand, being able to satisfy the cell’s energetic requirements but on the other, risking oxidative stress, appears to be valuable for interpreting our results. We note reasons for this robustness in Sections 3.4 and 3.6. Our results demonstrate that it is a combination of several factors that explains the differences in robustness observed in liver hepatocytes compared to the proximal tubule in the kidney, a tissue that is far less robust to ATP depletion. Our results indicate that the liver’s glycolytic activity, lower metabolic demand, and higher basline OXPHOS activites all contribute to the liver’s robustness to dysfunction of Complex III, the rate-limiting step of the electron transport chain, and all must be changed to PT-like values in order to produce PT-like sensitivity to OXPHOS dysfunction. Similarly we need to change all three of these features of the hepatocyte in order to see PT-like sensitivity to ATP depletion during hypoxia. The hepatocyte simply has more electron transport activity servicing a smaller metabolic demand with a larger contribution from glycolysis, and this makes it more robust to significant kinds of ATP depletion.

This ability to satisfy the cell’s energetic requirements may be examined in several ways. First of all, the cell is predicted to maintain a steady concentration of ATP even under a 50% increase in the maximal ATP consumption, as shown in Table 3. Second, as noted above in Section 3.4, if the rates of glycolysis are reduced we also see that the mitochondria can make up the difference by increasing their ATP generation. Each of these model simulations support our contention that the mitochondria may robustly provide for the hepatocyte’s ATP needs. These observations should not surprise us, since ATP depletion is not frequently observed in mitochondria-mediated liver pathologies, unlike oxidative stress [61, 41].

Ischemia-reperfusion injury is frequently caused by a temporary excessively reduced redox state of the electron transport chain causing oxidative stress during reperfusion [6]. We represent ischemia in our model by decreased oxygen tension and pyruvate concentration. During reperfusion we include a decreased supply of adenine nucleotides, based on a study that indicates a 75% smaller adenine nucleotide following ischemia [28]. We observe a transient redox state change as recorded in Figure 4. We also consider added effects from OX-PHOS dysfunction. When we include these effects, we see a more reduced coenzyme Q pool and a marginally lower transient proton motive force relative to the ischemia-reperfusion case without OXPHOS dysfunction, as shown in Figure 5. Both resemble experimentally observed timescales for the enhanced production of ROS following ischemia [11, 19, 26]. However the results in the latter case should be interpreted carefully. Reperfusion injury harms the electron transport chain, and it is often difficult to study the effects of ischemia on oxidative phosphorylation independently from the effects of reperfusion. For this reason, the levels of OXPHOS dysfunction used in Figure 5 should be treated as potential overestimates. These limitations are not a major problem because we are able to reproduce the appropriate timescale of reperfusion transients with or without OXPHOS dysfunction as noted above.

The saturation of the electron transport chain with electrons during reperfusion appears to be closely tied to the build up of succinate in ischemia, and the associated consumption of succinate following reperfusion. This coincides with suggested mechanisms for reperfusion injury in cardiac tissue [11]. As Chouchani et al. [11] notes, succinate build up has a preponderance of evidence indicating its importance to mitochondria-mediated ischemiareperfusion injury. Under more moderate hypoxia in other cases considered, where succinate build up is not observed in the same way, we do not see the same indicators of risk of ROS production that we note above.

In rats, it has been observed that chronic alcohol consumers have worse hepatic oxygen homeostasis relative to control non-alcoholic rats [47]. That is, in the event of acute alcohol consumption, chronic alcohol consumers may experience extremely low hepatic oxygen tensions. Sato et al. found that chronic alcohol consumers had a median oxygen tension upon acute alcohol consumption of 8 mmHg [47]. This is comparable to the hypoxic simulation case discussed in the previous paragraph. This suggests a mechanism by which alcohol consumption in individuals with alcoholic liver disease may trigger oxidative stress in a tissue, accelerating the progression of the disease.

While glycolysis may be elevated to levels like those noted previously in severe hyperglycemia, there are other ways to produce comparably elevated glycolysis as well. Fedatto et al. [15] studied hepatic tissue of rats with adjuvant arthritis (triggered by heat-killed *Mycobacterium tuberculosis*). The maximal rate of glycolysis was found to be roughly 30% higher in the arthritic rats. This presents another means of producing elevated glycolysis, which is potentially a trigger for oxidative stress. Fedatto et al. hypothesize other illnesses causing circulating inflammatory mediators may produce the same glycolytic effects [15], suggesting a range of diseases that could be causing oxidative stress in hepatocytes.

Our work can predict various indicators of oxidative stress, but cannot predict ROS production or the subsequent oxidative stress. Future work could our model with realistic models of ROS production. The former problem may be resolved by adapting existing models of ROS production [17] to the liver and incorporating them into our model. Predicting oxidative stress requires progress in the study of mitohormesis for the liver, so that thresholds for pathological ROS production may be found. As it stands we know that a moderate amount of ROS production may be a good thing [44]. Another contribution that could improve our model would be to integrate it with models of other pathways of hepatic metabolism, models of this kind exist for at least parts of hepatic metabolism [12]. Finally, our parameter estimates draw from a heterogeneous set of sources, including some experimental estimates from mice, namely, the Na-K-ATPase activity (discussed in Section 2.1), the cytosolic potassium concentration [57], and the pooled NADH/NAD^+^ concentration [8]. With more data from rats, it is possible we would have better predictions.

We include glutamate dehydrogenase activity in our model. Jonker et al. [27] found sexual dimorphism in the parameter K_NAD, ref_ in rat hepatocytes. For the purpose of this work an intermediate value was chosen between the sexes, but future work could expand on ours by creating a sex-specific version of the model that included this sexual dimorphism in glutamate dehydrogenase dynamics. Valle et al. [56] notes sexual dimorphism in OXPHOS capacity and mitochondrial protein content as well. These differences are likely to be the most impactful on mitochondrial behaviour since as we mentioned before, OXPHOS capacity is a key factor that explains the liver’s robustness to OXPHOS dysfunction.

Our results, obtained in this study and published studies, indicate that different tissues have characteristic ways that they are sensitive to mitochondrial dysfunction, or mitochondria-mediated injury. These ways can be connected to a range of mitochondrial and tissue-scale factors. For instance, the activity of OXPHOS enzymes is crucial to the robustness of the liver and is a mitochondrial-scale factor. On the other hand, tissue-scale factors such as high metabolic demand (necessary in the kidney for instance for urine concentration [1]) and oxygen tension appear in our model to be crucial for understanding the risks from mitochondrial diseases. This opens up space for extending our model: we know that the tissue may affect the mitochondria, but in aggregate, mitochondria may also affect the tissue. A limitation of our model is that tissue-scale factors are not modelled as dynamically responsive to mitochondrial functioning. For example, oxygen tension is clamped in our model, meaning that regardless of local oxygen consumption the tissue’s oxygen tension remains the same. Consequently, if we want to understand the effect of a certain degree of uncoupling (which increases oxygen consumption), we have no better method of pursuing this than simply seeing how it affects the mitochondria to combine that level of uncoupling with various degrees of hypoxia, without any way of predicting the appropriate degree of hypoxia. This limitation points towards a corresponding ambition however: multi-scale modelling of metabolism. We believe the best way to expand on this work is to integrate it into larger models of cellular transport [23, 31, 24, 22, 34, 36, 33, 35, 32, 38, 13, 30] and oxygenation [51, 50, 9, 10, 16] and of the factors determining local metabolic demand. Appropriate multi-scale models of metabolism will involve more than just placing our mitochondrial models inside of a larger scale (and ultimately, hopefully whole-body) model with appropriate interfaces. While the liver and kidney are particularly important to whole-body homeostasis, every tissue has mitochondria and most tissues are susceptible to mitochondrial dysfunction or mitochondria-mediated injury. For a whole-body multiscale model of metabolism then, we need models of mitochondrial respiration in other tissues.

In conclusion, we have developed a model of hepatocytes in mitochondria that predicts ATP generation, oxygen consumption, and risk factors of ROS production & oxidative stress. Our findings agree with empirical findings that ATP depletion is not a major risk in the liver, and seems to support the view that oxidative stress might be important to liver pathophysiology in many cases.

